# Characterization of ECM produced by MC3T3-E1 cellular spheroids encapsulated in alginate hydrogel

**DOI:** 10.1101/2025.01.15.633127

**Authors:** Diamante Boscaro, Lill Skovholt Wahlum, Marie Eline Ullevålseter, Berit Løkensgrad Strand, Pawel Sikorski

## Abstract

The application of cellular spheroids in bone tissue engineering research has gained significant interest in the last decade. Compared to monolayer cell cultures, the 3D architecture allows for more physiological cell-cell and cell-extracellular matrix (ECM) interactions that make cellular spheroids a suitable model system to investigate bone ECM in vitro. The use of 3D model systems requires fine-tuning of experimental methods used to study cell morphology, ECM deposition and mineralization, and cell-ECM interactions. In this study, we use MC3T3-E1 cellular spheroids encapsulated in an alginate hydrogel to study and characterize the deposited ECM. Spheroid shape and structure were evaluated using confocal microscopy. The deposited collagenous ECM was characterized using quantitative assay and microscopy, in particular Second Harmonic Imaging Microscopy (SHIM), hydroxyproline (HYP) assay and Transmission Electron Microscopy (TEM). The use of hydrogel constructs allows easy handling and imaging of the samples and helps to preserve the spheroid’s stability by preventing cells from adhering to the culture dish surface. We use a non-modified alginate hydrogel that does not facilitate cell attachment and therefore functions as an inert encapsulating scaffold. Constructs were cultured for up to 4 weeks. SHIM, HYP assay and TEM confirmed the deposition of collagenous matrix by the spheroid, with most of it taking place between week 2 and 4. We demonstrated that alginate encapsulated bone spheroids are a promising model for studying bone ECM in vitro.

## INTRODUCTION

To address bone-related diseases and injuries, bone tissue engineering (BTE) is currently focusing on the development of alternative cell-based methods that can overcome the limitation of traditional autologous and allogenic bone grafting (1, 2). These cell-based methods are relevant not only for regenerative applications, but also to improve the understanding of bone tissue biology. One of the most significant limitations of 2D cell cultures is the inability to reproduce the *in vivo* microenvironment. In the natural tissue, the cells are part of a complex 3D environment, in which the interactions with the surrounding cells and the extracellular matrix (ECM), together with the biological and mechanical stimuli, regulate cellular processes. For this reason, in the last decade, focus has been placed on the development of new cell-models that would be able to reproduce some aspects of the complex three-dimensional (3D) bone microenvironment found *in vivo*. To achieve this, several types of 3D *in vitro* cell models have been developed, such as organoids (3), organs-on-a-chip (4), and bioprinted tissues (5).

Another type of 3D cell culture is represented by the spheroid. Spheroids are characterized by a spherical shape, and form by spontaneous self-assembly of cells when cultured in a non-adherent environment (6). Spheroids are extensively used in multiple fields of biomedical research due to their ability to recapitulate aspects of natural tissues, such as the biological microenvironment (7, 8, 9), the cell-cell and cell-ECM interactions (10, 11), the soluble gradients (11), and the spatial morphology of natural tissues (12). In addition, they have shown apoptosis resistance (13, 14), secretion of anti-inflammatory molecules (14, 15), promoted cell viability, protein secretion and response to external stimuli (10, 11).

The ECM is a non-cellular, complex network of fibrous proteins and glycosaminoglycans with composition dependent on the type of tissue (16, 17). The ECM properties can influence cell-cell interactions, cell proliferation, and response to external stimuli (18). It maintains tissue stability and controls its mechanical properties, has a role in regulating the transport of molecules and soluble factors (19), as well as being a reservoir of growth factors and ions (20), and is responsible for the segregation between different tissues.

In bone, the ECM is involved in defining the functional properties of the tissue. Bone ECM is a complex and precisely structured inorganic-organic composite. The inorganic part consists of carbonate-substituted calcium-deficient apatite (20, 21), while the organic fraction consists of mainly type I collagen (90%) and proteoglycans, and non-collagenous proteins (10%), such as osteocalcin and osteopontin (22). Type I collagen is especially important since it helps in providing mechanical support and in controlling bone ECM mineralization (23). It has a triple-helix structure (known as tropocollagen molecules), consisting of three polypeptide chains of repeating Gly-X-Y units, where X and Y are usually proline and 4-hydroxyproline respectively (24). The tropocollagen molecules form fibrils (23) that can interact with other proteins found in the ECM (22). During bone mineralization, the mineral crystals will form in-between and on the surface of the collagen fibers (25). In addition, charged amino acids found in collagen bind non-collagenous proteins, which have a role in regulating mineral deposition (26).

Traditional methods used to investigate bone mineralization in monolayer cell cultures need to be adapted for proper characterization of this biological process in these 3D cell systems (3). Microscopy techniques, such as fluorescence or confocal microscopy, can face challenges due to possible limited diffusion of the dye (3) and/or attenuation problems in the aggregates (27). As infrared radiation (IR) is to a lower extent absorbed and scattered by biological tissues, IR microscopy techniques, such as two-photon fluorescence microscopy and Second Harmonic Imaging Microscopy (SHIM), are suitable for imaging of thicker samples (28), including hydrogel embedded spheroids. In addition, the possibility of doing a label-free imaging of deposited collagen matrix using SHIM, makes this method an advantageous technique when performing bone mineralization studies with these novel 3D cell models.

In this study, spheroids embedded in an alginate hydrogel were characterized and their ability to produce bone-like ECM was evaluated, with a focus on collagen deposition. Confocal microscopy was used to study cell morphology and aggregate structure in intact, embedded spheroids. We have evaluated the ability of the embedded spheroids to produce collagen matrix and the applicability of SHIM to image it. The results obtained were additionally supported by Transmission Electron Microscopy (TEM) images of the spheroids section. Overall, these results demonstrate that alginate-encapsulated spheroids are a promising model system for studying bone and bone mineralization *in vitro*.

## MATERIAL AND METHODS

### Cell culture

MC3T3-E1 subclone 4 (ATCC, CRL-2593) were cultured in tissue culture flasks in standard conditions at 37 °C and 5% CO_2_ in minimum essential alpha medium (MEM-α without ascorbic acid; ThermoFisher, A1049001) supplemented with 10% fetal bovine serum (FBS; Sigma-Aldrich). This media will be referred to as regular media (RM).

To induce osteogenic differentiation, RM was supplemented with 50 μg/ml (or 125.77 μM) of L-ascorbic acid 2-phosphate sesquimagnesium salt hydrate (Sigma-Aldrich, A8960) and 2 mM of β-glycerophosphate disodium salt pentahydrate (Sigma-Aldrich).

### Spheroids formation and diameter analysis

Spheroids were obtained using the micro-mold technique (29). Briefly, 1.5% agarose (Sigma-Aldrich) molds were made from #24-96 silicon molds (MicroTissues 3D Petri Dish micro-mold spheroids) (Figure 1 A and B) and were placed in a 24-well plate. The cells were collected and resuspended in 1 ml of RM, and 75 μl of cell suspension were added to each mold, followed by 1 ml of RM added into the well. After overnight incubation, the molds were placed upside down in the wells of the 24-well plate and centrifuged to allow spheroids collection. The molds were removed and the media with spheroids was collected in a 15 ml tube and centrifuged. The sedimented spheroids were retrieved and resuspended in a solution made of an equal volume of RM and sterile filtered 2% Alginate (Alg) solution (G fraction: 0.68, GG fraction: 0.57, Molecular weight: 250000 g/mol, [Novamatrix (Sandvika, Norway)]) (Figure 1 C). 150 μl of the spheroids solution was added into glassed-bottomed Petri Dishes (35 mm Dish with 14 mm bottom well, #1.5 glass – 0.16 - 0.19, Cellvis, Cat. #D35-10-1.5-N), covered with a GN-6 Metricel® 0.45 μm-47 mm sterile membrane and gelation was induced by adding 50 mM CaCl_2_ on top of the membrane for 5 minutes. After gelation, both the CaCl_2_ solution and the membrane were removed. The alginate disks containing spheroids were kept in 2 ml of media (Figure 1 D). Media was changed every second or third day. Spheroids were maintained with the same standard conditions as the cell cultures. Osteogenic media (OM) was used to induce osteogenic differentiation of the spheroids in the alginate disks.

**Figure 1:**
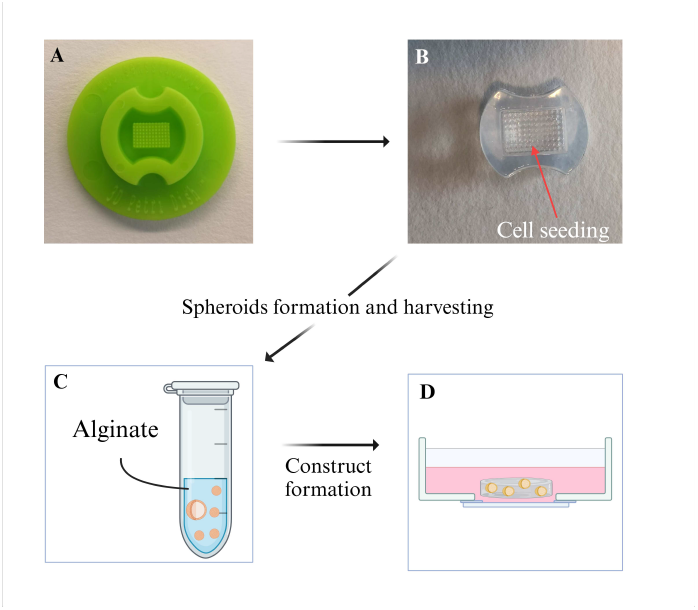
Schematic representation of spheroids formation. A) Silicon mold used to create the B) agarose molds. The cell suspension is placed in the central, pillared part of the mold. Spheroids are formed after ON incubation. C) After collection from the molds, spheroids are placed in a tube and resuspended in the alginate solution. D) After gelation in the glass-bottomed Petri dish, the alginate disk containing the spheroids is kept in 2 ml of RM or OM.

To assess the diameter, spheroids were measured on day 0 (encapsulation day), 3, and 7 using the Nikon ECLIPSE TS100 microscope. The diameter was measured using the software Fiji-ImageJ. The difference in diameter was compared between the RM and OM cultured spheroids.

### Morphological and structural characterization

To visualize the focal adhesions in spheroids after 3 weeks of culture, the immunostaining protocol from Weiswals et al. (30) was used. This protocol was used since it ensures the complete permeabilization of the spheroids, preservation of the structures, and localization of the protein of interest.

Briefly, samples from RM and OM cultures were washed with 5 mM BaCl for 5 minutes. Fixation and permeabilization were performed simultaneously, using 4% PFA and 1%Triton X-100 for 3 hours at 4 °C. The samples were washed three times for 10 minutes with PBS. Samples were dehydrated in ascending series of methanol (25%, 50%, 75%, 95%, 100%) at 4 °C in PBS and rehydrated following the same dilution series. Samples were then washed with PBS and blocked with 3% BSA + 0.1% Triton X-100 overnight at 4 °C. The following day, the samples were washed twice for 15 minutes, followed by incubation with primary antibody Paxillin (5H11) Monoclonal Antibody (Invitrogen, 1:200 in 1% BSA + 0.1% Triton X-100) for 48 hours at 4 °C. After incubation, the samples were washed four times for 30 minutes. Samples were incubated with Goat Anti-mouse IgG (H+L) Cross-Adsorbed secondary antibody Alexa Fluor^TM^ 488 (Invitrogen, 1:100 in 1% BSA + 0.1% Triton X-100) for 24 hours. The following day the samples were rinsed twice and stained for the nuclei with Hoechst 33258 (5 μg/ml) and for the cell membrane with CellMask^TM^ Deep Red plasma membrane stain (C10046) (Invitrogen, 1:1000 in PBS). The samples were then imaged using the Leica SP8 Confocal Microscope, with a HC PL Apo CS2 20x/0.75 water objective. Image analysis was performed using ImageJ.

To analyse the cell morphology, spheroids cultured in RM and OM for 2 weeks were fixed and permeabilized with 4% PFA and 1%Triton X-100 in Hepes buffer for 3 hours at 4 °C. The samples were then washed with Hepes buffer three times for 10 minutes and blocked with 3% BSA overnight. The cell actin was stained with Alexa Fluor 647 Phalloidin solution (Thermo Fisher; 1:100 in Hepes for OM samples and 1:200 in Hepes for RM samples), and cell nuclei were stained with Hoechst 34580 (Thermo Fisher, 1:1000 in Hepes), for 45 minutes at room temperature. Spheroids were imaged using Leica SP8 Confocal Microscope, with a HC PL Apo CS2 20x/0.75 water objective. Spheroids were imaged intact, embedded in the alginate hydrogel.

To evaluate if the reduced signal observed in the spheroids was caused by limited diffusion of the dye or by attenuation of the fluorescent signal, CellTracker Deep Red (ThermoFisher) was used to stain the cells before spheroid formation. Briefly, the cells in a tissue flask were stained with 3 μM solution of dye and RM, and incubated for 4 hours, before removing the staining solution and proceeding with spheroids formation, as described above. Imaging was performed using CLSM Zeiss 800 Airyscan Confocal Microscope, with a 10x/0.45 NA water objective.

### Second Harmonic Imaging Microscopy

Collagen production in monolayer cell cultures and alginate embedded spheroids was assessed using SHIM. Images were taken using a Leica SP8 SMD/MP Confocal Microscope, with a HCX IRAPO 25x/0.95 W (Leica Microsystems) water objective. The microscope was equipped with non-descanned detectors for detecting the SHG and two-photon excitation fluorescence signals. A 820 nm excitation was used, along with two emission filters: 390/40 nm for the Second Harmonic Generation signal and 445/20 nm for two-photon fluorescence.

Collagen production on monolayer cell cultures was assessed every week up to 4 weeks of culture in regular and osteogenic conditions. Cell were seeded in 6-well glassed bottom plates and were provided with RM and OM. At day 7, 14, 21, and 28 the cells were washed with PBS and stained for the nuclei using Hoechst 34580 (1:1000) for 20 minutes. After incubation, the cells were washed and imaged.

Collagen production in spheroids was assessed every week for up to 4 weeks of culture under regular and osteogenic conditions. Samples were imaged both unstained and stained for nuclei. For the stained samples, the gels were washed with PBS and stained using Hoechst 34580 (1:1000) incubated for 45 minutes. After incubation, the gels were washed and imaged.

The images were processed using ImageJ, and the background subtracted to remove the signal overlap between the two channels.

Due to the partial overlap of the emission wavelengths of the collagen (410 nm) and the nuclei dye, which causes some attenuation of the signal coming from the collagen, we decided to perform imaging of unstained and stained spheroids sample. The two-photon fluorescence emission cube was also used during the imaging of the unstained samples. This overlap and attenuation effect was observed both in monolayer cell cultures samples and spheroids.

### Transmission electron microscopy

For TEM analysis, 1, 2 and 3 weeks old RM and OM cultured alginate gels with the embedded spheroids were washed with 1 ml of 5 mM BaCl, followed by fixation with 4% formaldehyde and 2.5% glutaraldehyde in PBS at 4 °C for 24 hours. The fixative was removed, and the samples were cut into 1-2 mm pieces and placed in an eppendorf tube, covered with PBS for storage. For preparation, the samples were rinsed three times in 0.1 M sodium cacodylate for 10 minutes, followed by post-fixation in 1% osmium tetroxide for 1 hour. The samples were then rinsed three times for 10 minutes in water, followed by staining with 2% uranyl acetate in water for 1 hour. Dehydration was performed in increasing acetone series (25 %, 50%, 75%, 90%, 96%, and 2×100%) for 10 minutes each. The samples were then embedded in a pure acetone and Epon resin mix in a ratio of 2:1, 1:1 and 1:2 for 30 minutes each. The samples were then incubated in Epon resin, which was changes three times at intervals of 1 hour, overnight and 1 hour respectively. The samples were embedded in fresh resin and polymerized at 60 °C for two days. The samples were sectioned using a Leica EM UC6 ultra-microtome at 60 nm and collected on copper TEM grids with formvar support film. Ultra-thin sections were post-stained with Reynold’s lead citrate. Spheroids sections were imaged using a FEI Tecnai 12 with a tungsten filament, with an acceleration voltage of 80 kV.

### Hydroxyproline assay

Hydroxyproline (HYP) assay was performed to assess collagen quantity in monolayer cell cultures and embedded spheroids, cultured in RM and OM. The data obtained from the HYP assay were normalized to the DNA content for each respective sample.

For 2D cell cultures, collagen quantification was performed after 1, 2, and 4 weeks of cultured in RM or OM. On the day of the analysis, the cells were washed with PBS, resuspended, and centrifuged. The cell pellets were resuspended in 200 μl of ultra-pure water. 100 μl were used to perform the HYP assay, while the remaining was used to perform the DNA quantification.

For spheroids, collagen quantification was performed after 1, 2, and 4 weeks of cultured in RM or OM. To collect the embedded spheroids, the hydrogels were moved to a 1.5 ml Eppendorf tube and dissolved by adding 50 mM of citrate solution, and vortexed for about 5 to 10 minutes, until gel dissolution. The gels were dissolved together to avoid sample loss. The tubes were then centrifuged at 1000 rpm for 5 minutes. The supernatant was removed and the pellet was trypsinized. After resuspension in media, the samples were centrifuged, collected, and resuspended in 200 μl of ultrapure water. 100 μl were used to perform the HYP assay, while the remaining was used to perform the DNA quantification. The HYP assay was performed as described in (31, 32). Briefly, 100 μl of cell suspension/spheroids suspension was transferred to a pressure-tight propylene vial and 100 μl of HCl (38%) were added. The vial was capped with a Nalgene TM PPCO Low-profile Cap and was placed in the oven at 110 °C for 18 hours for to allow hydrolysis of collagen. The vials were centrifuged at 10000 x g for 3 minutes and dried in the oven at 50 °C (48/72 hours). After drying, samples were resuspended in 200 μl of ultra-pure water and vortexed briefly. 60 μl of each sample in triplicate were transferred to a 96 well plate. Trans-4-hydroxy-L-proline was used as standard. To each sample and standard, 20 μl of Assay Buffer (composed of 1-propanol, ultra pure water and citrate stock buffer. See supplementary info for more details) and 40 μl of Chloramine T reagent were added. The plate was then covered with aluminum foil and incubated for 20 minutes at room temperature. After the addition of 80 μl of DMBA reagent, the 96 well plate was placed in the over at 60 °C for 60 minutes. The plate was left to cool down and the absorbance at 570 nm was measured using a the SpectraMax i3x plate reader (Molecular Devices).

DNA content was measured using the PicoGreen Quant-iT Assay Kit (Invitrogen). Cells were lysed in 0.2% Triton X-100 solution and shaken for 30 minutes before the DNA content measurement.

### Statistical analysis

Data from monolayer cell culture represent triplicates derived from three independent samples. Data from spheroids represent triplicates derived from a minimum of (independent) 2 alginate disks containing each 10 to 25 spheroids. Data are shown as mean value ± standard deviation. Statistical analysis was performed using a oneway ANOVA with a Turkey’s post-hoc test. P < 0.05 was considered significant. Significantly different data points are denoted with the same symbol when obtained from different time-points and with an * when comparing data from the same time-point. Weak-significant (0.05 < P < 0.15) and non-significant difference were noted with *ws* and *ns* respectively.

## Results

### Characterization of the embedded spheroids: size, shape, morphology, and interaction with the surrounding hydrogel

The aim of this study is to create and characterize alginate encapsulated spheroids with a focus on longtime culture, compatibility with microscopy and other techniques that can be used to study the produced ECM and consequently evaluate their osteogenic potential. In our model, spheroids are randomly distributed within an alginate disk that is approximately 1 mm in thickness and 5 mm in diameter, composed of 1% alginate hydrogel. The construct is maintained in a glass-bottomed Petri dish, supplied with RM or OM. The osteogenic potential of OM cultured spheroids is evaluated and compared to spheroids cultured in RM and monolayer cell cultures. Alginate was chosen for ita ECM-like properties and its widespread use in the field of TE (33). In this study, alginate hydrogel is used as an inert scaffold that allows culture and characterization of encapsulated spheroids over a prolonged culture period. Alginate hydrogel prevents cells form attaching to the culture dish surface, while alginate discs are easy to handle during the culturing period, and are compatible with a variety of microscopy techniques. Spheroids from MC3T3-E1 cell line were obtained using the micro-mold technique using agarose molds (29). The micro-mold technique was selected for spheroids production as it allows easy handling of the samples during the procedure, the possibility of obtaining a relatively high number of spheroids per batch, good control over the number of cells per spheroid, and has good reproducibility between the different batches. Spheroids in an agarose mold before harvesting are shown in Figure 2. Due to cell-cell interactions and the self-assembly process, spheroids compact during the first days after preparation. This also happens when they are encapsulated in the alginate hydrogel. Spheroids size was reduced by almost 50% between day 0 and day 7, with an average diameter for respectively RM and OM of 230.79 ± 24.25 μm and 231.03 ± 33.87 μm on day 0 (encapsulation day), 145.06 ± 22.66 μm and 132.79 ± 21.28 μm on day 3 and 113.14 ± 19.12 μm and 109.93 ± 20.86 μm on day 7 (Figure 3 A to G). Significant changes in size were not observed after day 7.

**Figure 2:**
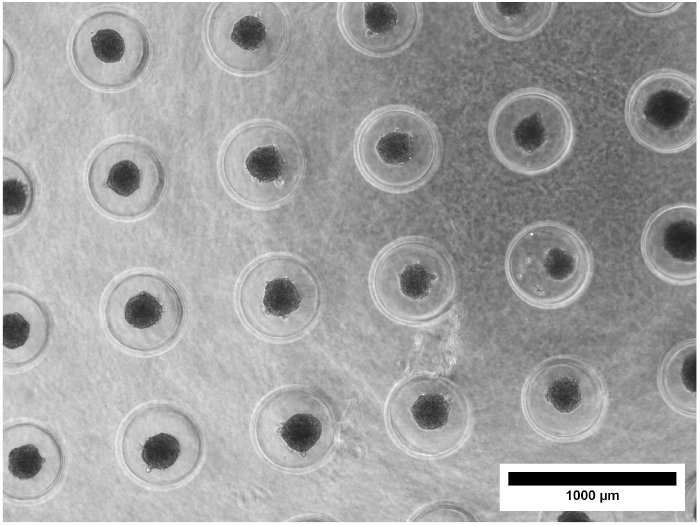
Spheroids (gray spheres) in the mold, obtained after overnight incubation.

**Figure 3:**
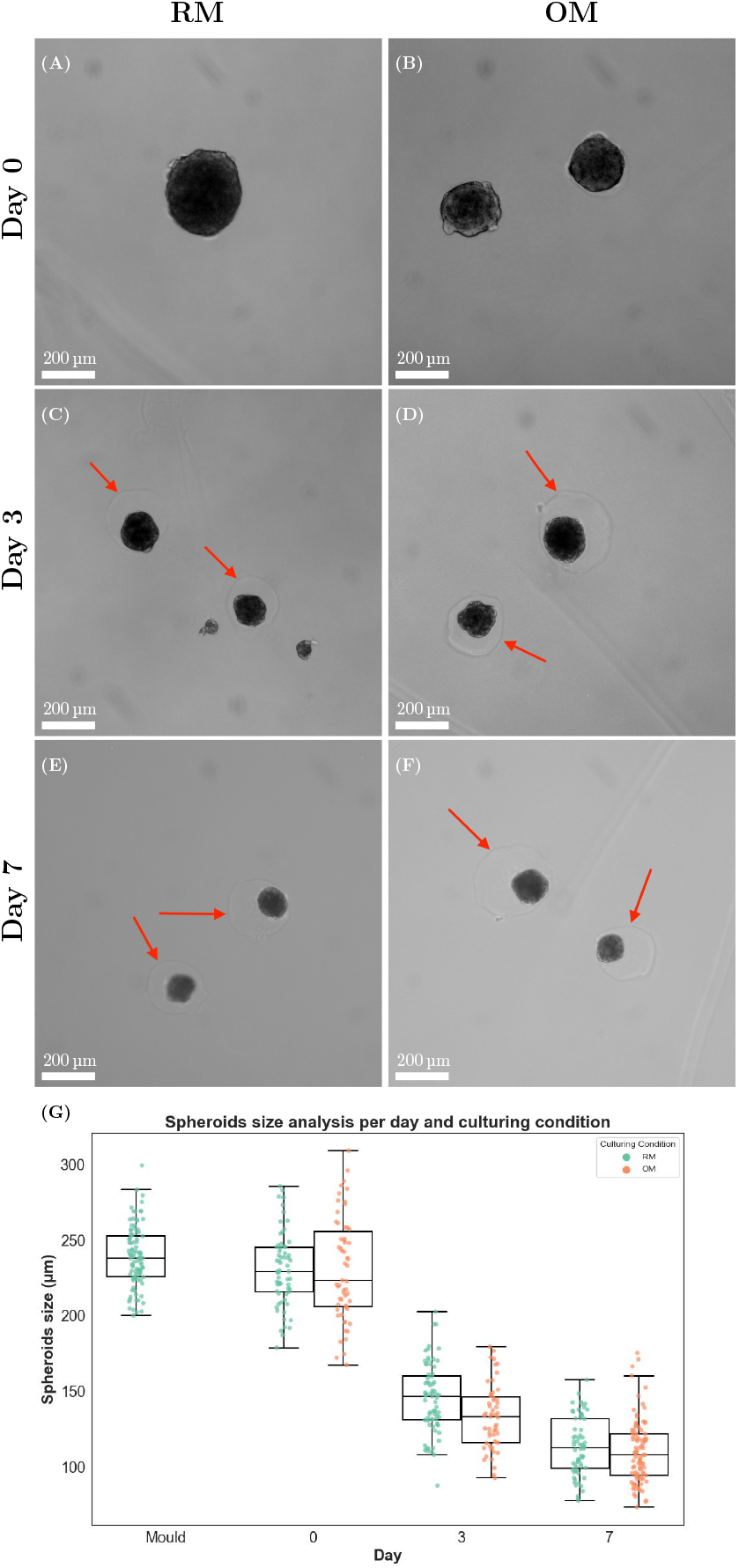
Spheroids reduce their size during the first week of culture. (A-F) Bright-field microscopy images of the spheroids encapsulated in the alginate hydrogel. The red arrows indicate the outline of the pocket that is formed due to the size reduction. (G) Quantification of the diameter reduction of spheroids cultured in RM and OM during the first week: spheroids were measured in the mold after overnight incubation (n= 97), after encapsulation (day 0) in RM (n=64) and OM (n=58), day 3 in RM (n=73) and OM (n=55) and day 7 in RM (n=64) and OM (n=96).

As the spheroid compact, a small pocket is formed in the alginate hydrogel (red arrow in Figure 3 C to F). The size of this pocket is consistent with the size of the spheroids at the encapsulation time-point (Day 0). In spheroids cultured for more than 1 week, it was noticed the presence of small particles in the pocket area, usually close to the edges of the pocket, in both RM and OM cultured spheroids. These structures were identified using transmission electron microscopy (TEM) as extracellular vesicles and cellular debries (FigureS1).

The shape of the spheroid was also monitored during the culturing period. The shape was retained during the first two weeks of culture in both RM and OM. Around the third week of culture, it was observed that around half of the spheroids cultured in RM began to show loss of spherical shape and the presence of protrusions (Figure 4 upper left). Immunostaining for focal adhesions’ Paxillin of these samples revealed heavily stained structures in the protrusion area (Figure 4 lower left, right side of the dotted white line). Major structural changes were not observed in OM cultured spheroids, which retained their spherical shape (Figure 4 upper and lower right). Figure S2 shows how these structures were also stained using a membrane dye, revealing the presence of lipidic structures in the protruding area.

**Figure 4:**
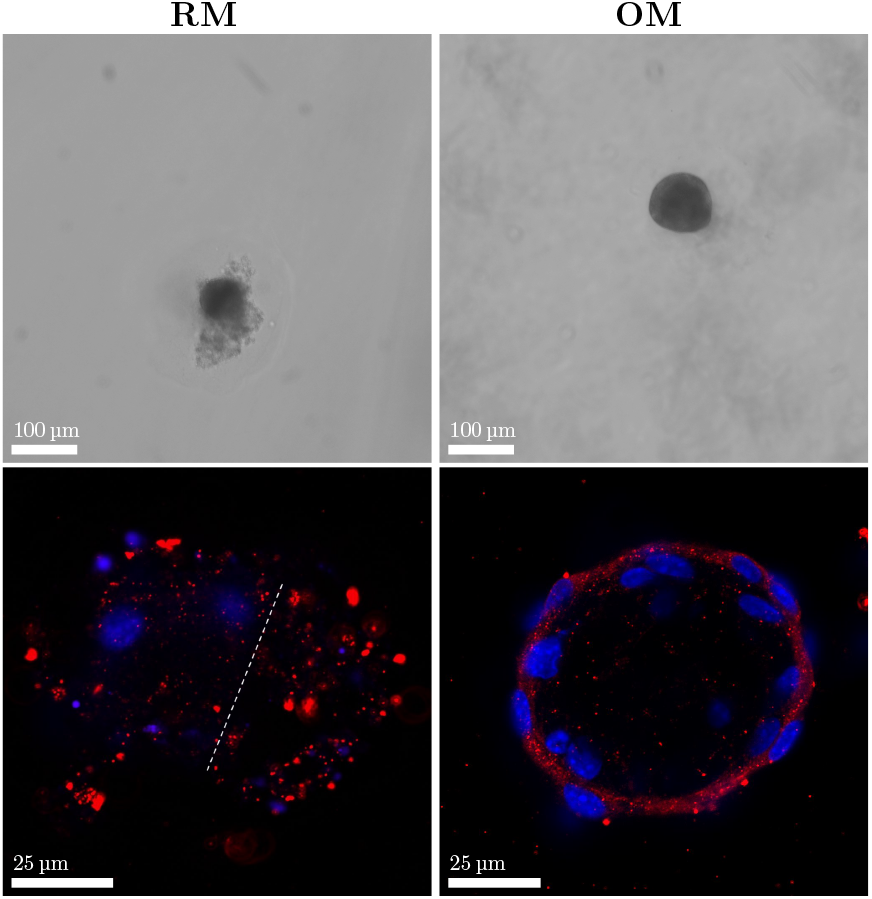
Shape analysis of 3 weeks RM and OM cultured spheroids. Upper images show bright field images of the spheroids, and it can be appreciated the difference in shape between the two different cultured samples. Lower images show immunofluorescence images of spheroids stained with nuclei (blue) and focal adhesion (red).

To assess the response to the differentiation stimuli, we analyzed and compared the morphology of the cells in spheroids cultured in RM and OM. When grown in a tissue flask, MC3T3-E1 cells in the pre-osteoblasts stage are characterized by a rounder, more spread morphology, but when cultured in differentiating conditions, they show a more elongated shape (Figure S3). To assess any change in the morphology of the cells in the aggregate, spheroids cultured in RM and OM for 2 weeks were stained for the nuclei and actin filaments. Spheroids grown in RM displayed round nuclei and an overall round cell structure, as it can be observed by the stained actin filaments (Figure 5). This round structure was not observed in the OM cultured spheroids, which displayed a thinner, more elongated structure with a more compacted nuclei shape in some cells in the outer layers (Figure 5).

**Figure 5:**
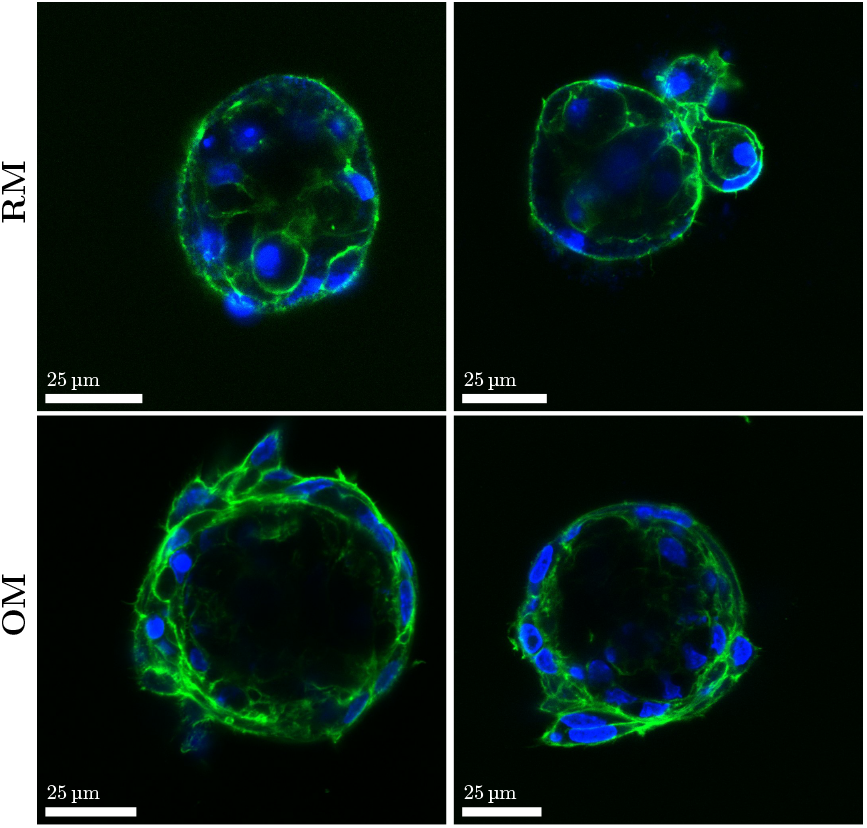
Cell morphology is influenced by the culturing conditions. Fluorescence images of actin (green) and cell nuclei (blue) of spheroids after 2 weeks in RM and OM.

During confocal microscopy imaging, a weaker signal was observed inside the spheroids. This can result from a restricted dye diffusion or an attenuation of the fluorescence signal. To verify the cause, we stained cells with CellTracker Deep Red before spheroid formation, and then investigated the signal intensity as a function of the position. The data obtained indicated that the loss of signal is caused by the signal attenuation, as observed in images obtained at different positions along the z-axis in the same spheroid (Figure 6 and Figure S4). This result is consistent with 2-photon fluorescence microscopy described below, where a more uniform signal from the whole aggregate was detected.

**Figure 6:**
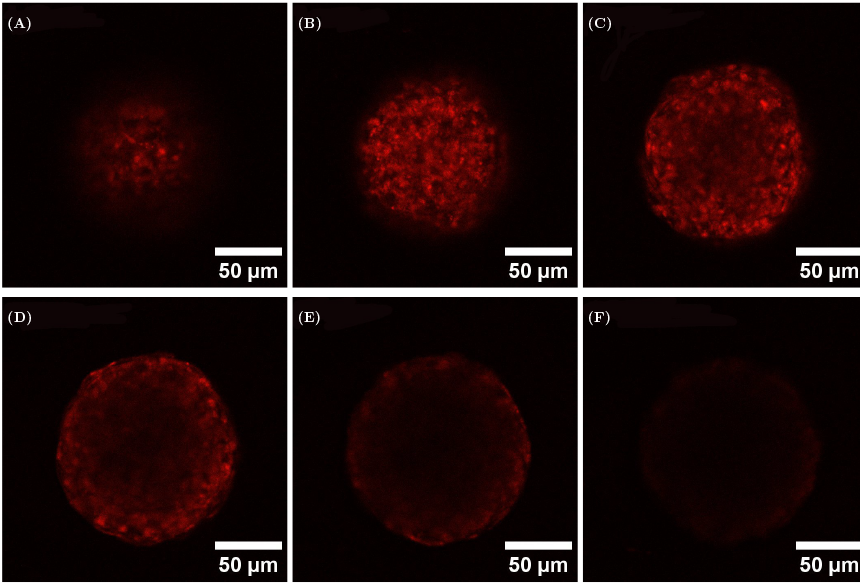
Analysis of optical effects on spheroids, treated prior to aggregation with CellTracker Deep Red. The 6 cross-sections were taken with a step of 20 μm between each other, with the first section (A) starting at 5 μm.

### Osteogenic potential of encapsulated spheroids

The potential of the encapsulated spheroids to produce a bone-like matrix was then investigated, with focus on the production of collagen. Collagen deposition in spheroids in OM and RM media was assessed every week for up to 4 weeks in a qualitative manner using SHIM and quantitatively using HYP assay. SHIM was performed on stained and unstained samples (Figure 7 A and S5). To confirm that the observed signal was originating from the deposited collagen, SHIM was performed also on monolayer cell cultures (Figure S6). The first detectable collagen deposition using SHIM was observed in spheroids cultured in OM for 14 days (Figure 7A). A progressive signal increase can be observed in week 3 and 4 OM cultured spheroids, corresponding to an increase in collagen deposition by the cells in the aggregates (Figure 7 A), with no signal detected for RM cultured samples at any of the time-points. Collagen deposition was limited to the spheroids volume, with no collagen observed in the pocket area and surrounding gel. TEM imaging of spheroids confirmed the presence of collagen fibers in samples cultured for 2 and 3 weeks in OM, but also revealed the presence of collagen in samples cultured for 1 week in OM (Figure 8, region in the red rectangle). No collagen was detected in TEM images for RM cultured spheroids (Figure 8).

**Figure 7:**
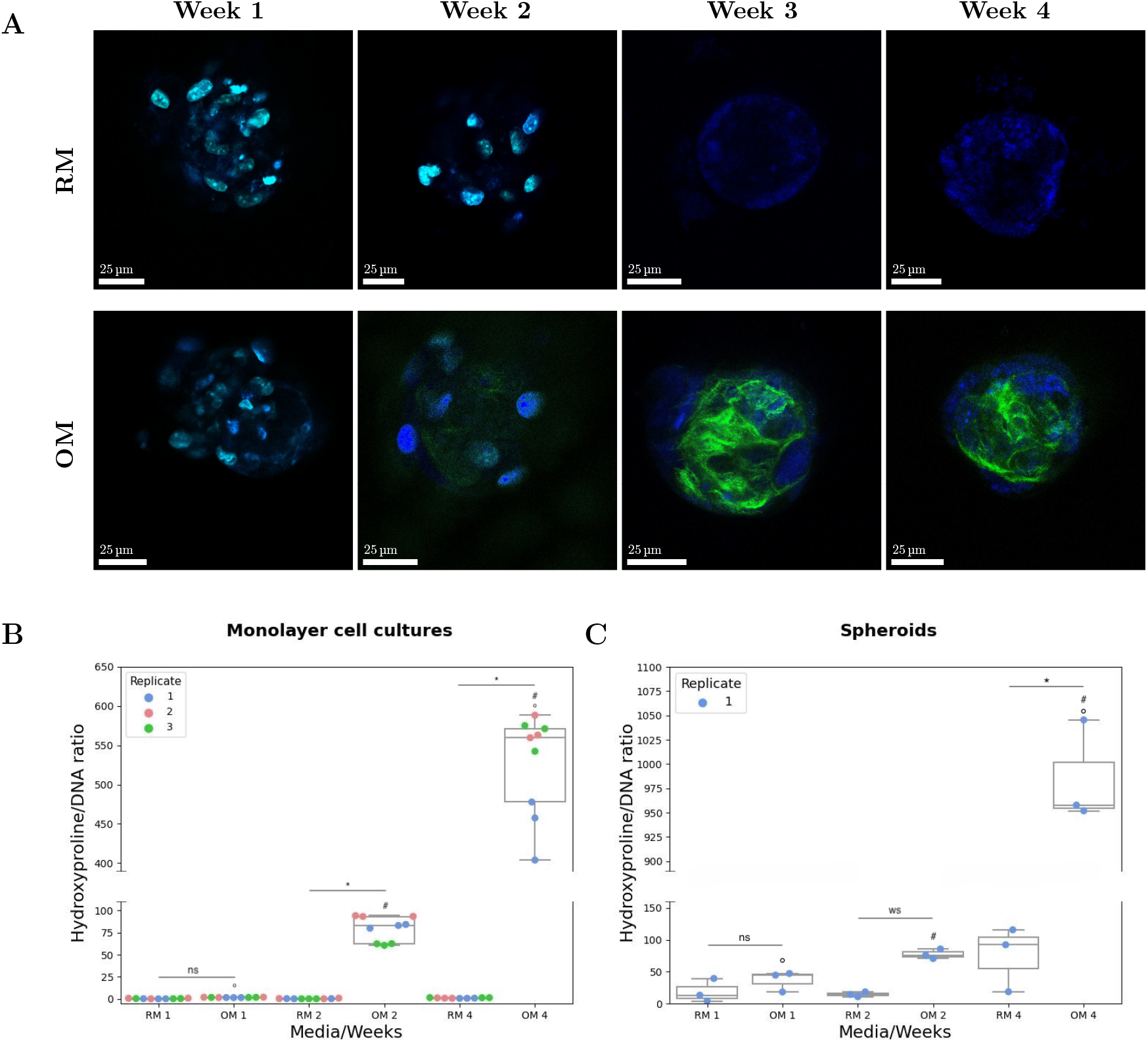
Osteogenic potential of alginate encapsulated bone spheroids. A) SHIM images of RM and OM cultured spheroids over 4 weeks. Collagen deposition (green) can be observed in 2, 3 and 4 weeks OM cultured spheroids. Spheroids cultured for 1 and 2 week were stained with Hoechst 34580 (blue), while spheroids cultured for 3 and 4 weeks were left unstained to ensure better collagen signal observation. Biochemical quantification of collagen in B) monolayer cell cultures (n=3) and C) spheroids (n. of disks=2). The HYP data were normalized to the DNA content. In spheroids, weak-significant difference was observed between RM and OM cultured samples for week 2 (p = 0.121) and statistical difference was observed between RM and OM cultured samples for 4 (p < 0.001), while for week 1 the data were considered non-significant (p = 0.979).

**Figure 8:**
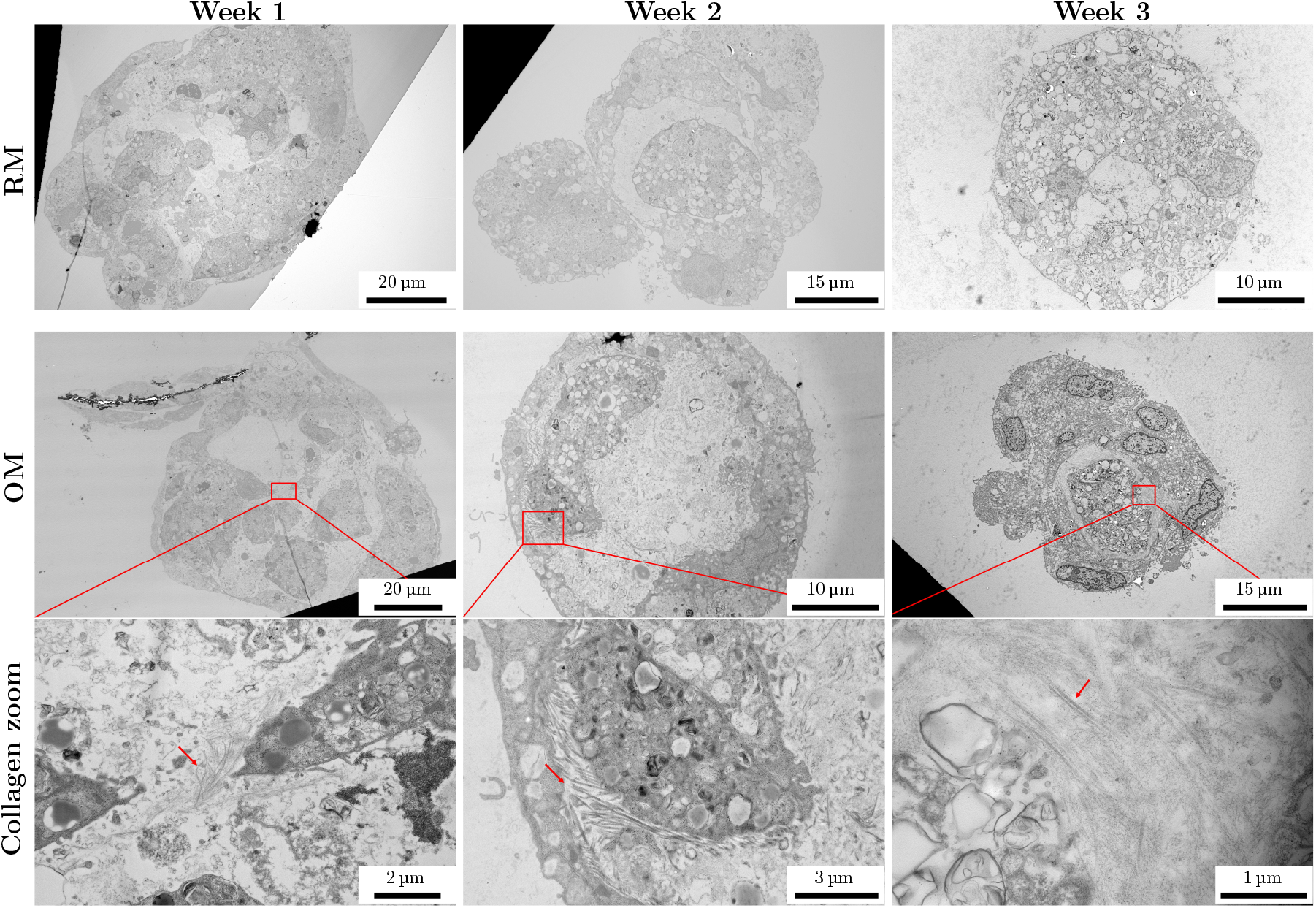
TEM images of 1, 2 and 3 weeks old spheroids cultured in RM and OM. Regions of OM spheroids with collagen fibers are noted with a red rectangle and the zoomed region is shown below. Collagen fibers are pointed with the red arrow.

Biochemical quantification of HYP confirmed the results obtained with both imaging techniques. Specifically, the HYP assay was consistent with the TEM results for 1 week OM cultured spheroids, in detecting the small quantity of collagen produced by the cellular aggregates, despite the statical analysis indicating no significant difference between RM and OM at week 1. The biochemical quantification revealed also that at the final time point (4 weeks), more collagen had been deposited by the spheroids compared to the monolayer cell cultures (Figure 7 B-C).

## Discussion

3D cell models, such as spheroids, have been shown to better mimic certain aspects of natural tissues, making them a good cell model to study tissue biology (34). In addition, the development of 3D models that can support bone regeneration is interesting from the bone tissue engineering perspective. For this reason, proper characterization of the deposited ECM and mineral phase and evaluation of the ability of these models to reproduce the natural cellular environment are required since they influence the properties of developed tissue. In this work, we characterize bone spheroids encapsulated in alginate hydrogels. We focus on the production of collagen matrix, as it is the first step in bone tissue formation and a prerequisite for bone mineralization studies. Our long term goal is to investigate the ECM mineralization in these aggregates and to characterize the deposited mineral. In this work, we demonstrated that culturing spheroids in hydrogel discs can be advantageous for handling a large number of samples and for performing microscopy. Spheroids were made using MC3T3-E1 subclone 4 murine cells. This cell line was used as they are known to be a suitable model for investigation of the ECM and its mineralization (35). As extensively described in literature (36, 37), this cell line is easy to grow, and differentiation into osteoblast-like cells can be induced by treatment with ascorbic acid and β-glycerophosphate. To perform more complex bone tissue development studies, a system that shows osteocytic differentiation is required, aspect that lacks when using the MC3T3-E1 cell line (35). Application of MC3T3-E1 cell line is advantageous when performing short term analysis of differentiation and optimizing techniques used for characterization, for example of the produced ECM.

We observed that spheroids in alginate were stable and viable, and that the alginate helped in maintaining the culture for a prolonged period. This is required for the analysis of the ECM deposition and for further analysis of the ECM mineralization process. TEM confirmed that the cells were not able to attach to, nor invade the alginate matrix, providing further insight into the environment created in the pocket and the deposited ECM and collagen by the spheroids.

In OM cultured spheroids, deposition of the collagenous matrix was detected using both quantitative assay and microscopy. The presence of the alginate hydrogel allows for easy handling and localization of the sample while performing microscopy analysis. The presence of the hydrogel did not influence the imaging process. Due to the nature of our sample, we decided to use SHIM as a non-invasive, label-free technique to monitor collagen deposition. To our knowledge, this method has not been used previously for the analysis of collagen in intact spheroids embedded in a hydrogel. The results obtained using this technique are in line with already published results where collagen analysis was carried out using traditional techniques (38, 39). With these results, we demonstrated that SHIM can be used as an alternative microscopy technique for collagen imaging. SHIM allows for imaging of the whole aggregate, including the inner regions; an aspect that lacks when using confocal microscopy. In addition, due to the nature of the technique, the samples do not require excessive handling and preparation, limiting the possibility of sample damage. The possibility of analyzing living samples is also advantageous in terms of reusing the samples for other types of analysis and for limiting contact with dan-gerous substances that are usually used for fixing purposes. We consider the attenuation effect to be specific to our case. The SHG signal obtained from the collagen can be generated from different values of excitation wavelengths, which traditionally spaces from 700 to 1000 nm (40). For our specimens, the optimal excitation value resulted in a partial overlap of the emission wavelength with the emission of the nuclear dye. Nonetheless, for the purpose of our investigation, this was not considered to be a major limitation.

Implementation of the quantitative assay is required for sample collection prior to the analysis. Using a citrate solution to dissolve the alginate hydrogel proved to be an efficient method for sample collection, with minimal sample loss. For spheroids samples, the amount of cells is relatively low, and a sensitive assay to detect produced collagen quantitatively is needed. The HYP assay used in this work has a sufficient sensitivity to detect collagen produced by approximately 30 spheroids at each time point and type of culture, where each spheroid contains around 3000 to 3500 cells. We observed that the deposited collagenous ECM had a role in supporting the stability and the shape of the spheroid. After culturing the spheroids for around 3 weeks, we observed that RM cultured spheroids started to show a loss of the spherical structure, with presence of lipidic structures, ECM proteins, and cells in these less organized region. OM cultured spheroids, on the other hand, did not show any significant changes. At this time point, OM cultured spheroids contain a significant amount of collagen, which we believe is contributing to the stability of the cell aggregates. The observed protrusion in RM cultured spheroids can originate from the cells that are potentially detaching or dividing from the outer layer. The focal adhesion immunostaining performed on the RM cultured samples revealed the presence of stained structures in the protrusions. Due to the absence of the collagen matrix, these structures are not confined to the surface and are able to grow outside of the aggregate. These results suggest that the cells in the outer layer may have the ability to proliferate to some extent, although the topic of cell proliferation within the aggregate remains under debate (11, 13, 41).

## Conclusions

We demonstrated that alginate encapsulated bone spheroids are a promising model to study bone mineralization in vitro. SHIM for collagen imaging in intact spheroids was found to be a highly effective technique for monitoring collagen matrix deposition, allowing for in-depth imaging of the aggregate, which is not achievable with traditional imaging techniques.

## ACKNOWLEDGEMENTS

This project was founded by the Norwegian University of Science and Technology (NTNU). TEM sample preparation, after the fixation, was performed in the Laboratory of Electron Microscopy of the Nencki Institute, supported by the project financed by the Minister of Education and Science based on contract No 2022/WK/05 (Polish Euro-BioImaging Node “Advanced Light Microscopy Node Poland”). We also acknowledge Stiftelsen Biopolymer for financial support.

