## Supplementary info for "Characterization of ECM produced by MC3T3-E1 cellular spheroids encapsulated in alginate hydrogel"

### Supplementary information for: Characterization of ECM produced by MC3T3-E1 cellular spheroids encapsulated in alginate hydrogel

**Diamante Boscaro**[a,✉], **Lill Skovholt Wahlum**[a,✉], **Marie Eline Ullevålseter**[a,✉], **Berit Løkensgrad Strand**[b,✉], **and Pawel Sikorski**[a,✉]

[a]Department of Physics, Faculty of Natural Sciences, Norwegian University of Science and Technology (NTNU), Trondheim, Norway
[b]Department of Biotechnology and Food Science, Faculty of Natural Sciences, Norwegian University of Science and Technology (NTNU), Trondheim, Norway

#### A. MATERIALS AND METHODS.

***A.1. HYP assay.*** The assay buffer is composed of 1-propanol, ultrapure water and citrate stock buffer to a volume ratio of 3:2:10. The citrate stock buffer is composed of citric acid anhydrous ($4.61\,\mathrm{g}$), anhydrous sodium acetate ($7.22\,\mathrm{g}$), sodium hydroxide ($3.4\,\mathrm{g}$), and acetic acid ($1.26\,\mathrm{mL}$), with a volume of $100\,\mathrm{mL}$ and pH 6.

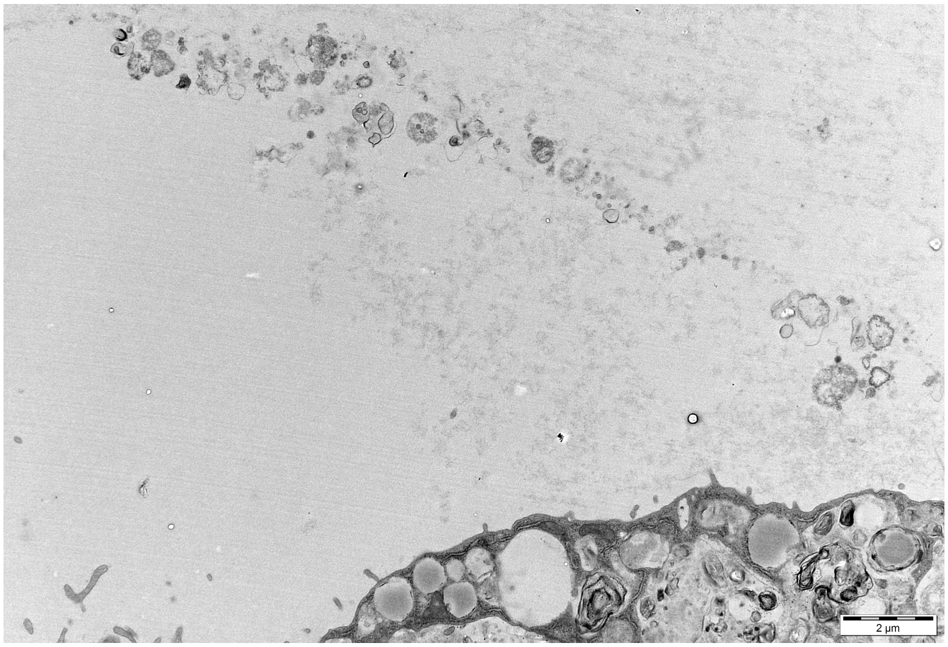


Figure S1: TEM image of the vesicles released in the pocket. The edge of the spheroid can be seen at the bottom of the figure, while the extracellular vesicles and the cellular debries can be seen as the small, darker, spherical structures. These vesicles are usually observed to be closer to the edge of the pocket. Their presence show and confirm that no interaction is happening between the spheroid and the hydrogel.

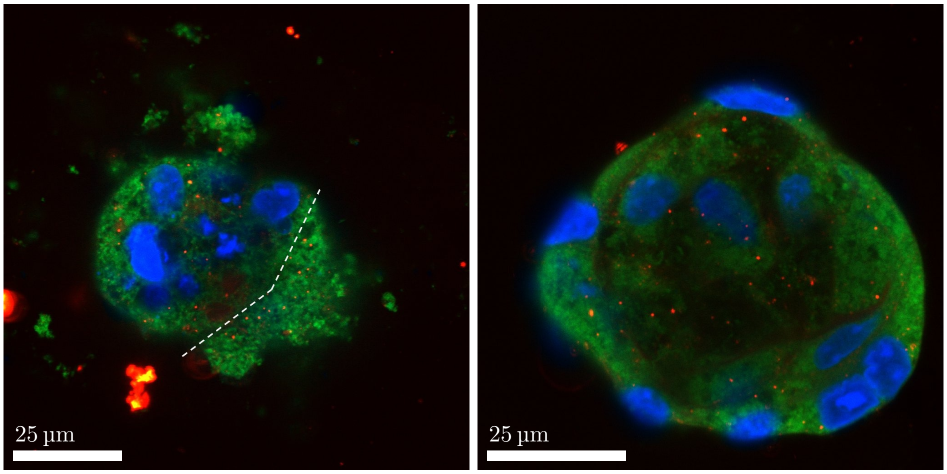


Figure S2: Shape analysis of 3 weeks RM (left) and OM (right) cultured spheroids. Nuclei are shown in blue, focal adhesions in red and cell membranes in green. It can be noticed that the protrusion area in the RM cultured spheroids (right side of the dotted white line) was stained for both focal adhesions and cell membrane.

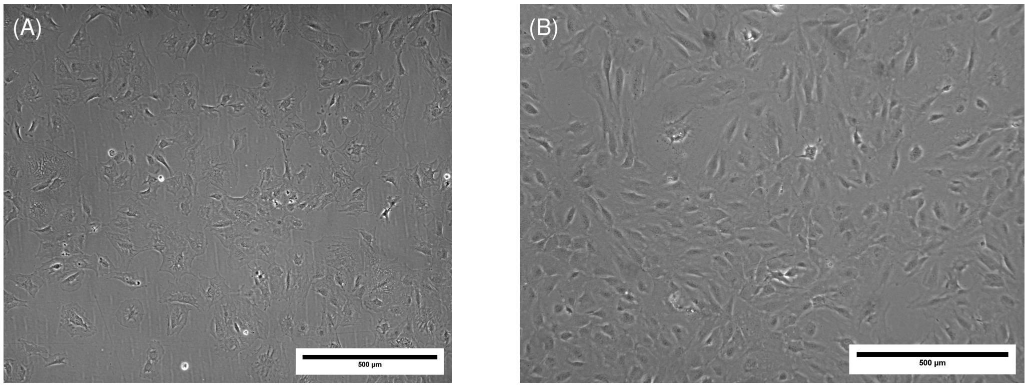


Figure S3: Evaluation of cell morphology in monolayer cells cultured in RM and OM. Figure A shows cells cultured in RM and Figure B shows cells cultured in OM. Cells seeded in RM show a more spread shape compared to OM cultured cells, characterized by a more elongated morphology. The same characteristic was observed in cells making up the spheroid.

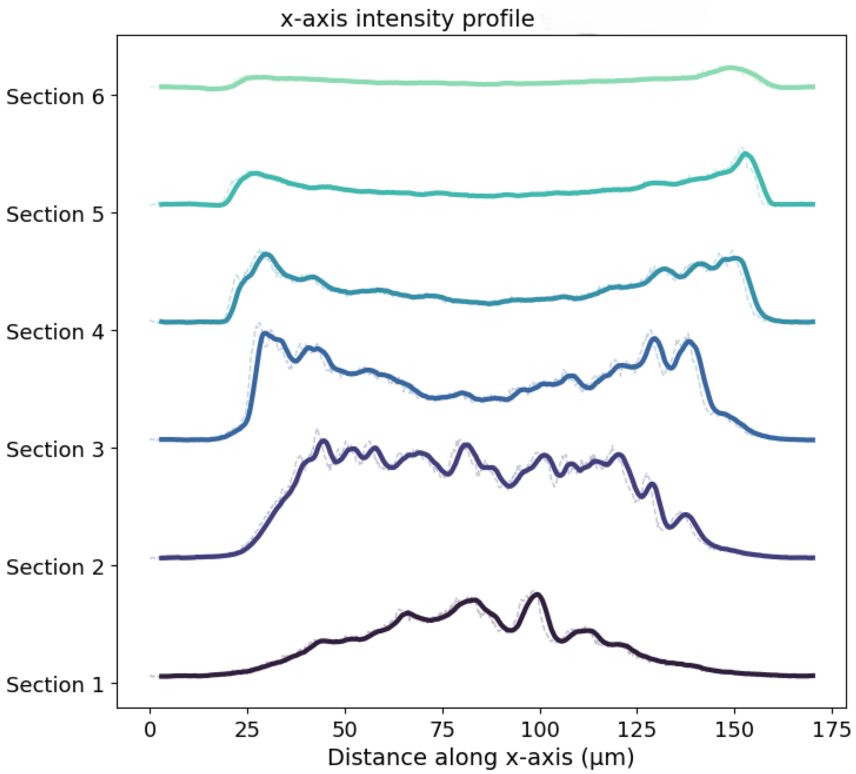


Figure S4: Intensity profile collected from RM cultured spheroid, 2 days after encapsulation, using CLSM. Results were plotted as intensity variations as a function of the distance along the positive x-axis, where x is the diameter of the spheroid along the x-axis ($138\,\mu m$). The interval between each section is $20\,\mu m$, with a total Z-stack height of $135\,\mu m$. The data demonstrates that the fluorescence signal is significantly attenuated by the dense cell aggregate. Original data points are shown as dashed lines and solid lines represent interpolated data.

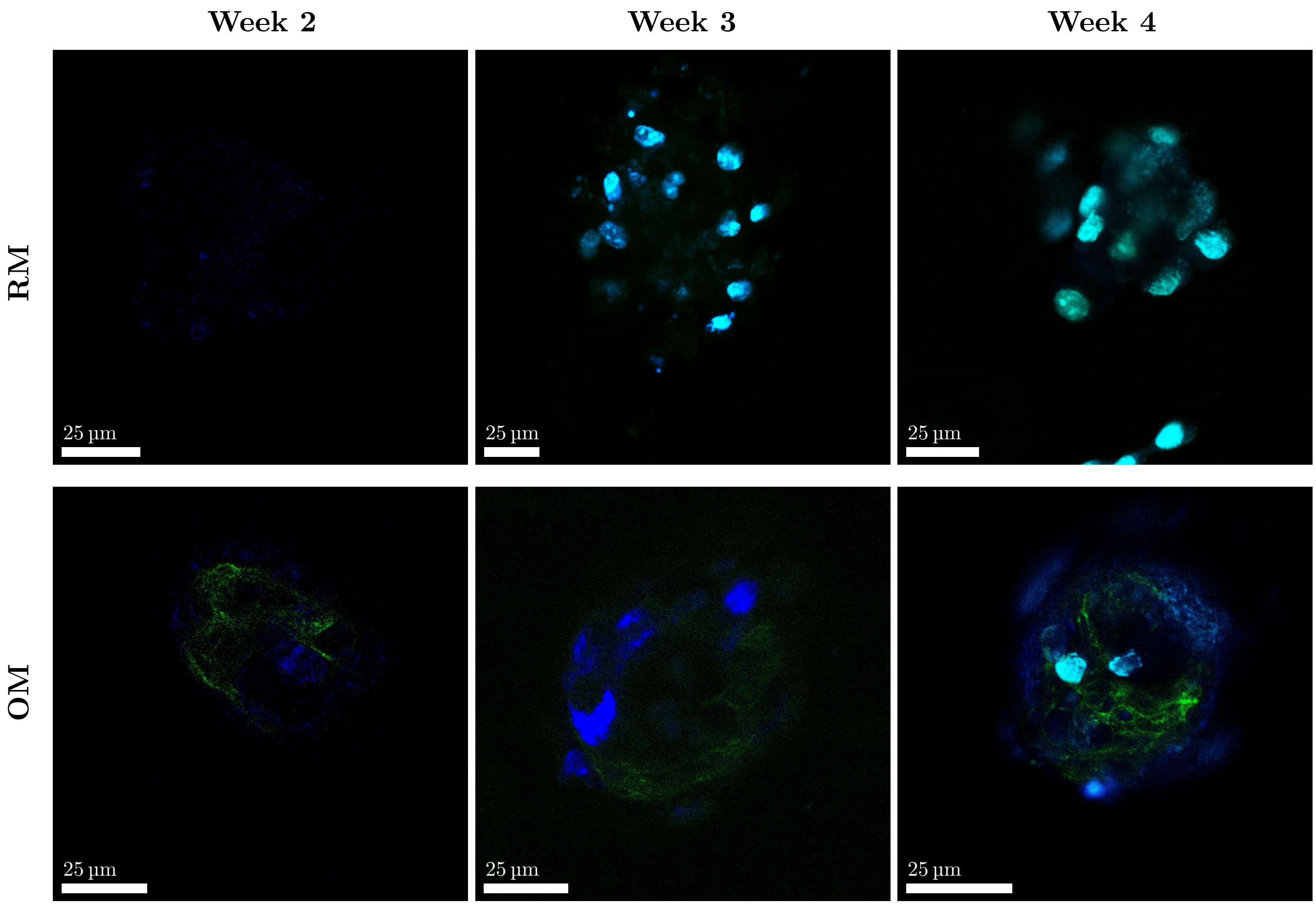


Figure S5: Collagen deposition in spheroids. 2 weeks old samples were left unstained for the nuclei, while 3 and 4 weeks old samples were stained for the nuclei (blue). The blue signal coming from the unstained samples resulted to be auto-fluorescence from the cells. Collagen fibers can be seen in green. A strong signal attenuation can be observed when comparing stained and unstained samples at the same time point.

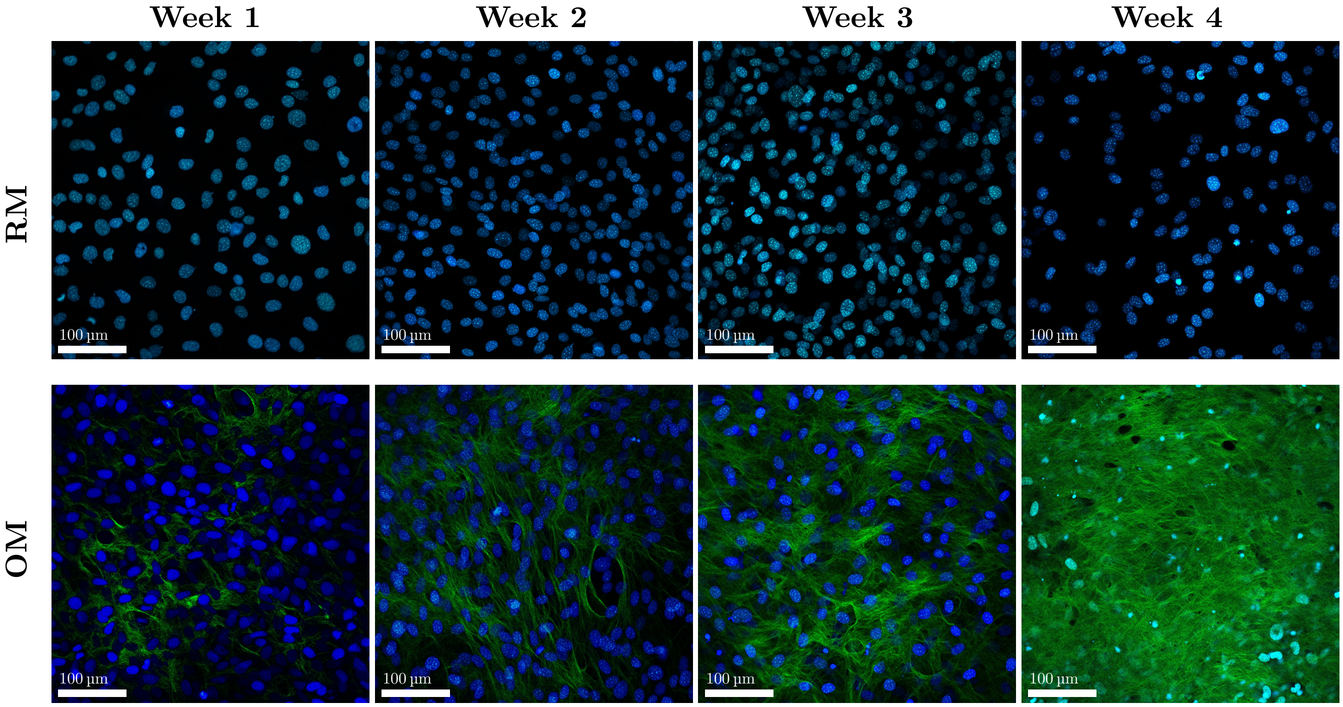


Figure S6: Collagen deposition in monolayer cell cultures. Cells were cultured in RM and OM for 4 weeks and were imaged for collagen production every week up to 4 weeks. The images show nuclei (blue) and collagen (green). A progressive increase in the presence of collagen can be observed starting from week 1, where small and localized spots of deposited collagen were observed, to week 4, where a uniform layer of collagen lays on top of the cells. No collagen deposition was observed in RM cultured samples.
